## Supplemental data for "Inclusive, Exclusive and Hierarchical Atlas of NFATc1^+^/PDGFR-α^+^ Cells in Dental and Periodontal Mesenchyme"

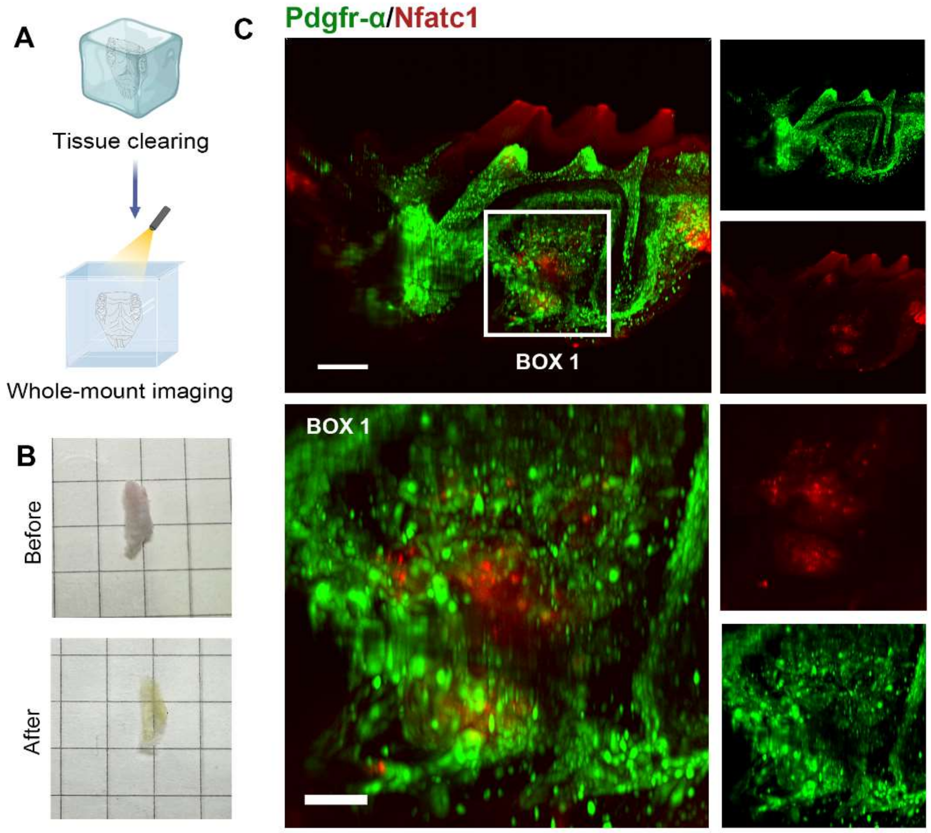


**Figure S1.** A. The tissue-clearing (TC) and whole-mount imaging procedure of mice maxilla. B. The images before & after the TC procedure of mice maxilla. C. The distribution of PDGFR-a$^{+}$ and NFATc1$^{+}$ cells from the section of XZ axis after 3D reconstruction of TC imaging. The image was from *Pdgfr-α$^{CreER}$; Nfatc1$^{DreER}$; LGRT* mice (pulse). Box 1: alveolar bone. Scale bar = 300 μm for top row, 100 μm for bottom row.

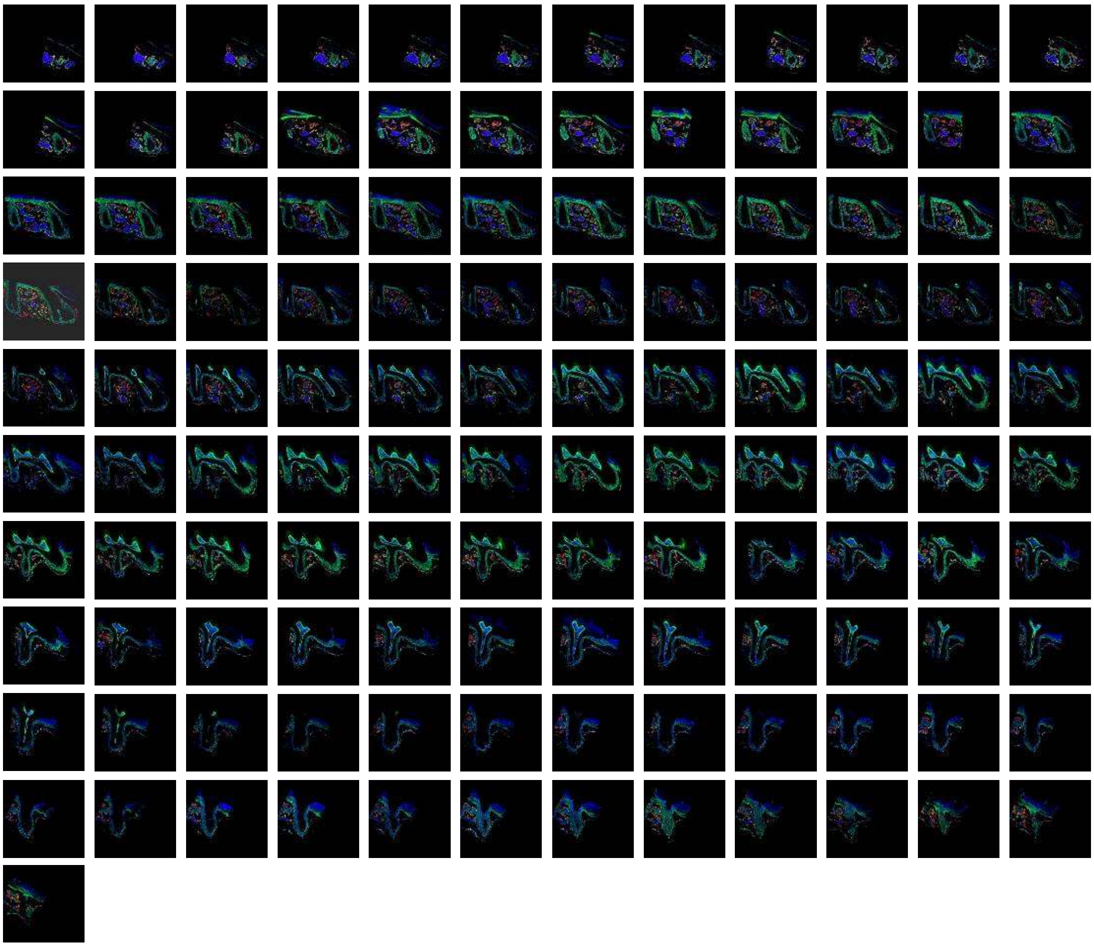

**Figure S2.** The total 121 slices of maxilla M1 of *Pdgfr-α*$^{CreER}$*; Nfatc1*$^{DreER}$*; LGRT* mice sample (pulse). The images were acquired by confocal microscope, ZsGreen$^{+}$ cells in green, tdTomato$^{+}$ cells in red, DAPI in blue.

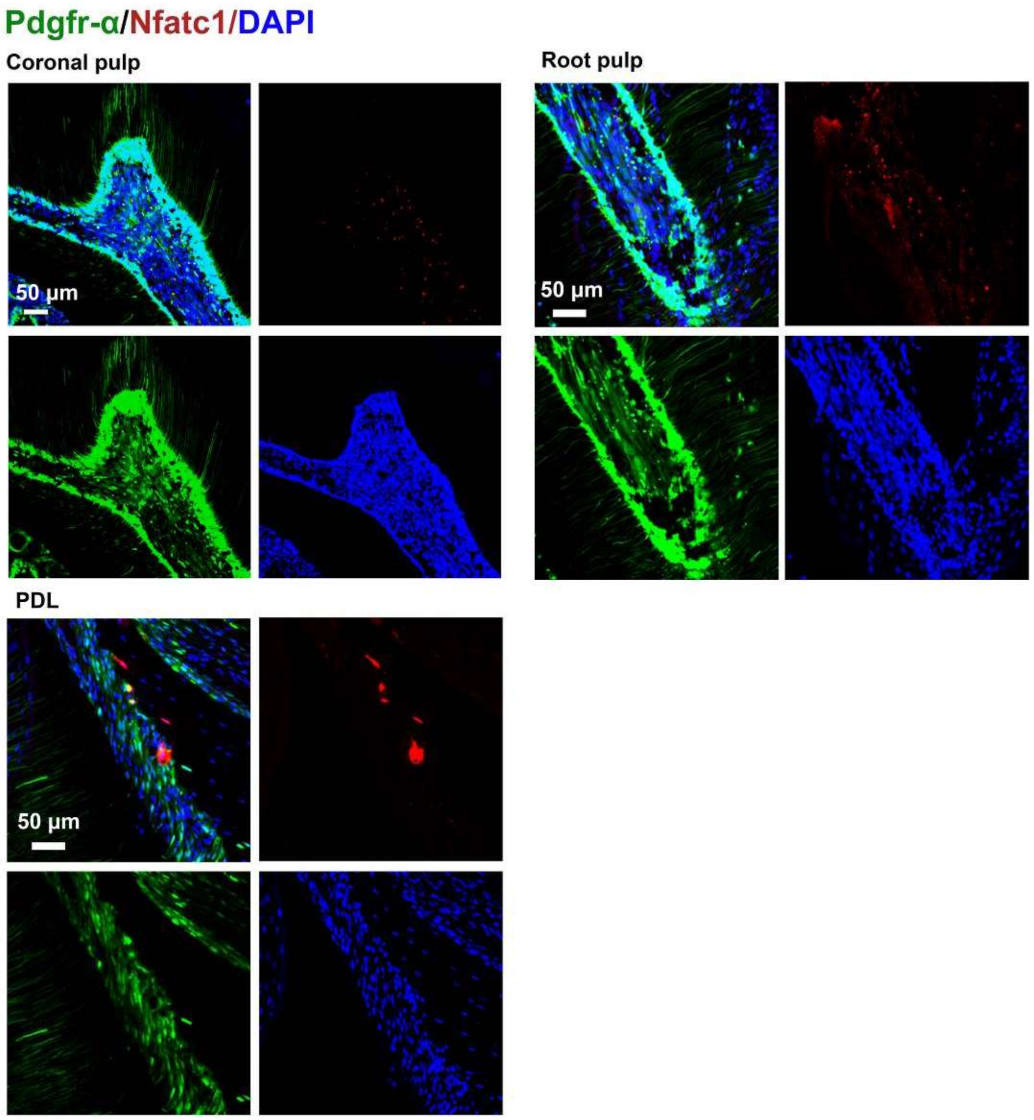


**Figure S3.** Representative images of coronal pulp, root pulp, and PDL of maxilla M1 of *Pdgfr-α*$^{CreER}$*; Nfatc1*$^{DreER}$*; LGRT* mice sample (pulse). The images were acquired by confocal microscope.

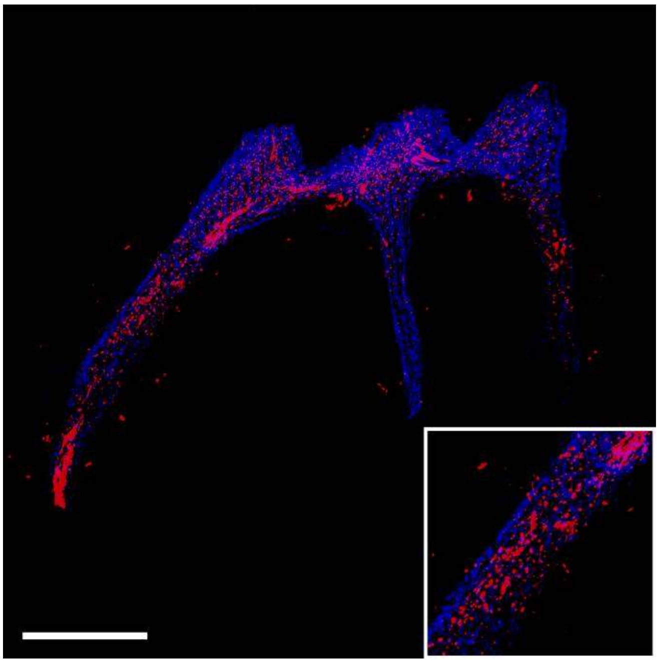

**Figure S4.** The tdTomato signal in pulp reconstructed by traditional serial section-based confocal imaging method (scale bar = 300 μm). The sample was from *Pdgfr-α*$^{CreER}$*; Nfatc1*$^{DreER}$*; LGRT* mice (pulse).

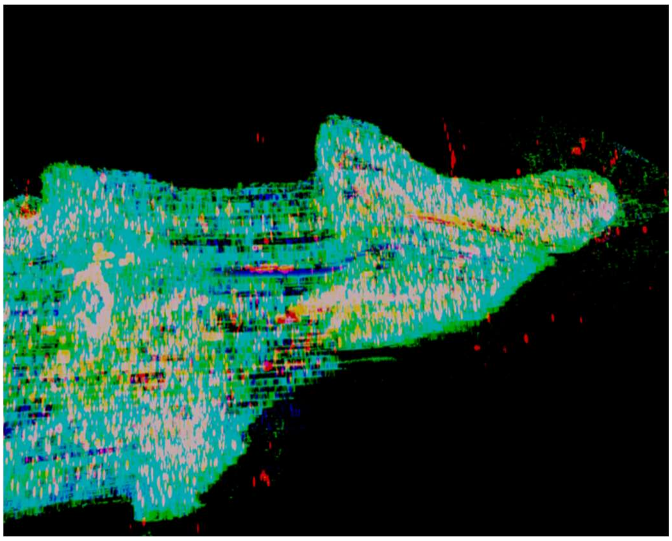

**Figure S5.** The discontinuities in the z-axis due to stratification of slices. The image was reconstructed by serial sections of maxilla M1 of *Pdgfr-α*$^{CreER}$*; Nfatc1*$^{DreER}$*; LGRT* mice sample (pulse).

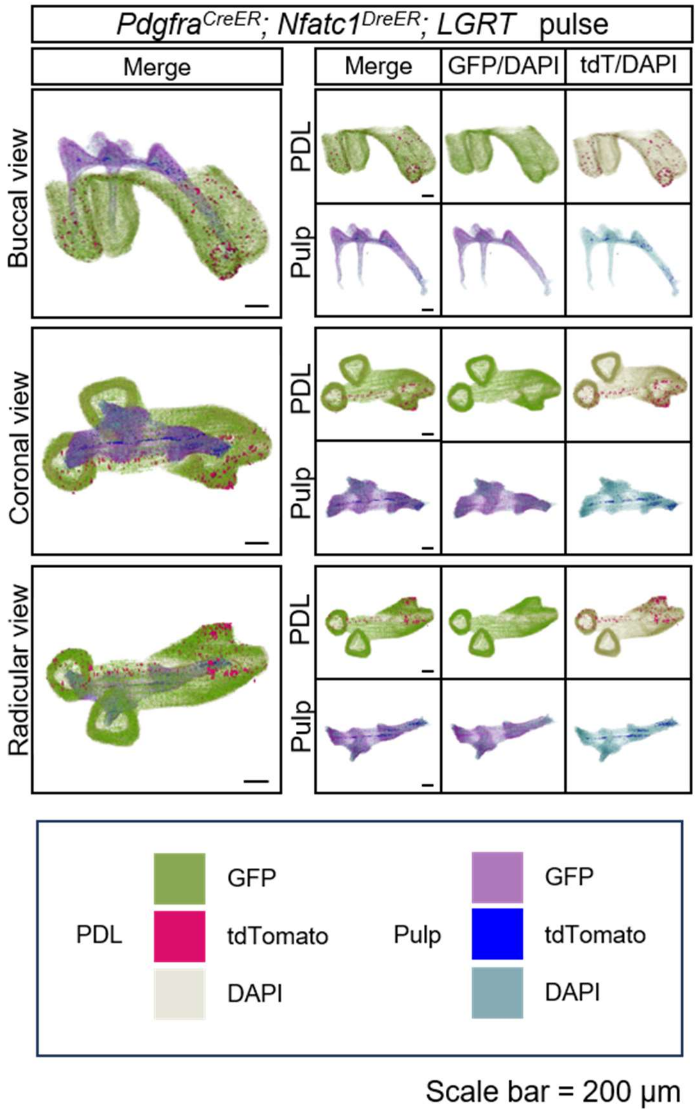


**Figure S6.** 3D reconstruction of maxilla M1 of *Pdgfr-α*$^{CreER}$*; Nfatc1*$^{DreER}$*; LGRT* mice (pulse) by DICOM-3D; in PDL: ZsGreen$^{+}$ cells in green, tdTomato$^{+}$ cells in rose red; in pulp: ZsGreen$^{+}$ cells in purple, tdTomato$^{+}$ cells in blue. The image stack was also displayed in buccal view, coronal view, and radicular view of pulp and PDL, respectively.

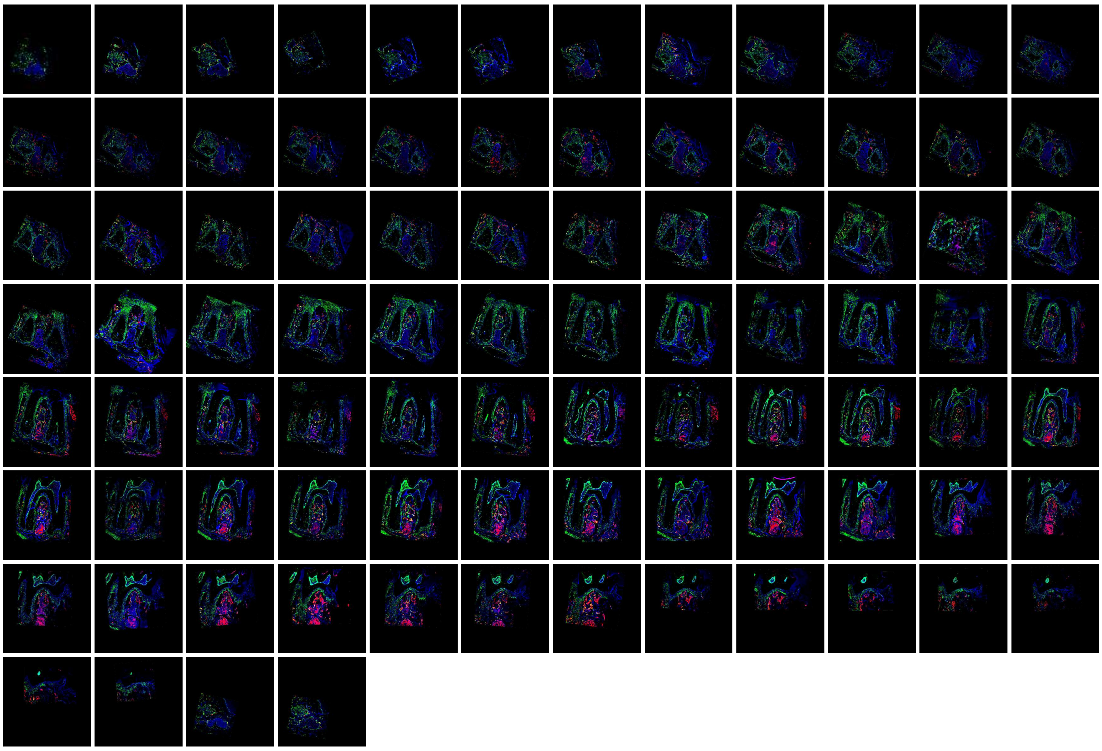

**Figure S7.** The total 88 slices of mandible M1 of *Pdgfr-α*$^{CreER}$*; Nfatc1*$^{DreER}$*; LGRT* mice sample (pulse). The images were acquired by confocal microscope, ZsGreen$^{+}$ cells in green, tdTomato$^{+}$ cells in red, DAPI in blue.

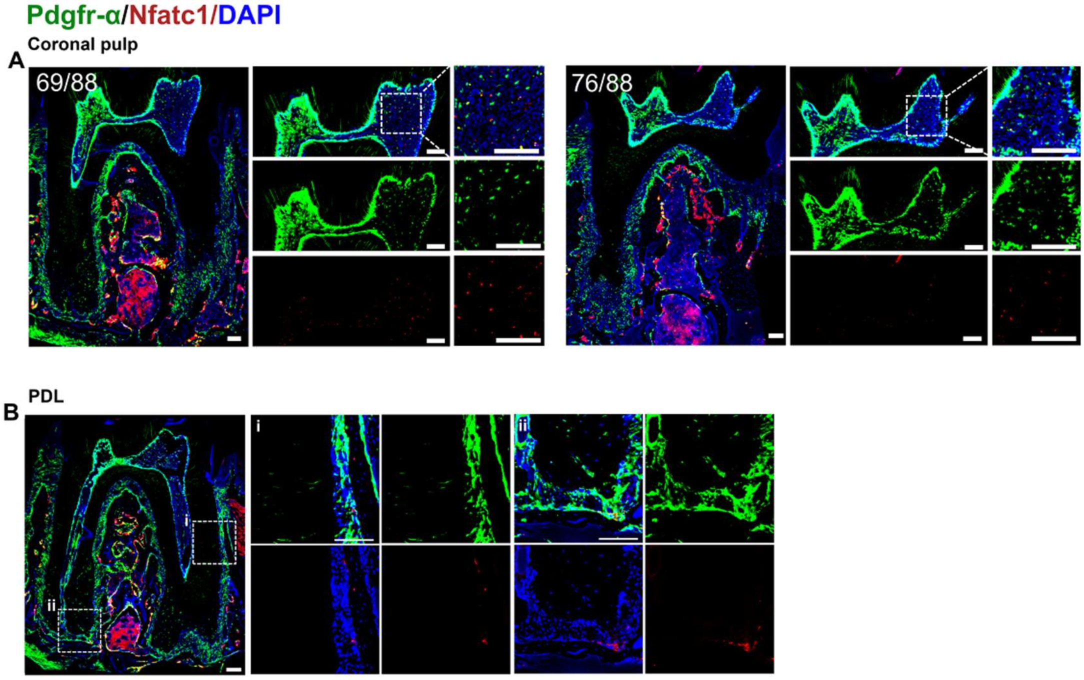


**Figure S8**. Representative images of coronal pulp (A) and PDL (B) acquired by confocal microscope of mandible M1 of *Pdgfr-α*$^{CreER}$*; Nfatc1*$^{DreER}$*; LGRT* mice sample (pulse). Scale bar = 100 μm

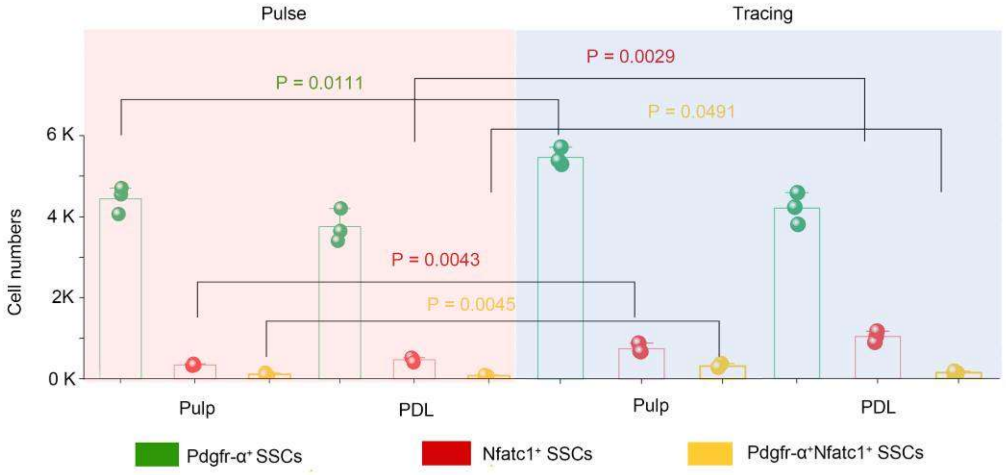


**Figure S9.** The number of PDGFR-α$^{+}$ cells, NFATc1$^{+}$ cells, and PDGFR-α$^{+}$&NFATc1$^{+}$ cells in pulp and PDL, respectively. The quantification was done on a z stack of images in Imaris using the automatic spot detection feature.

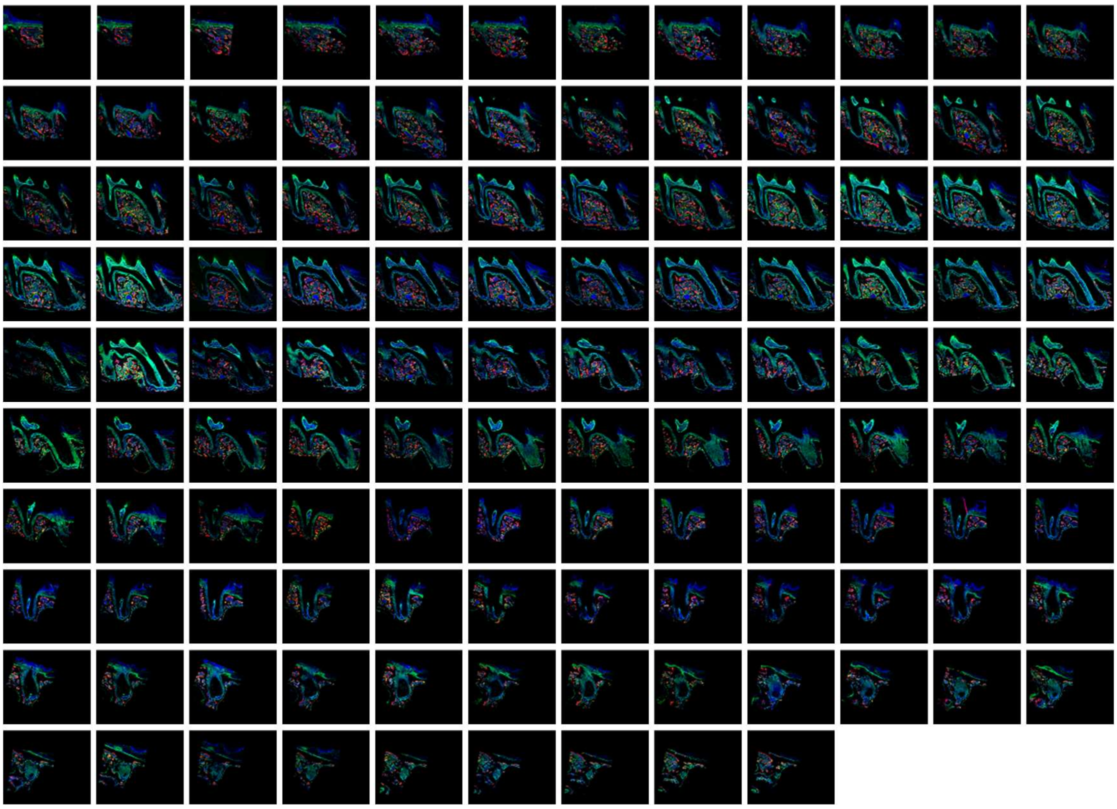

**Figure S10.** The total 117 slices of maxilla M1 of *Pdgfr-α*$^{CreER}$; *Nfatc1*$^{DreER}$; *LGRT* mice sample (tracing). The images were acquired by confocal microscope, ZsGreen$^{+}$ cells in green, tdTomato$^{+}$ cells in red, DAPI in blue.

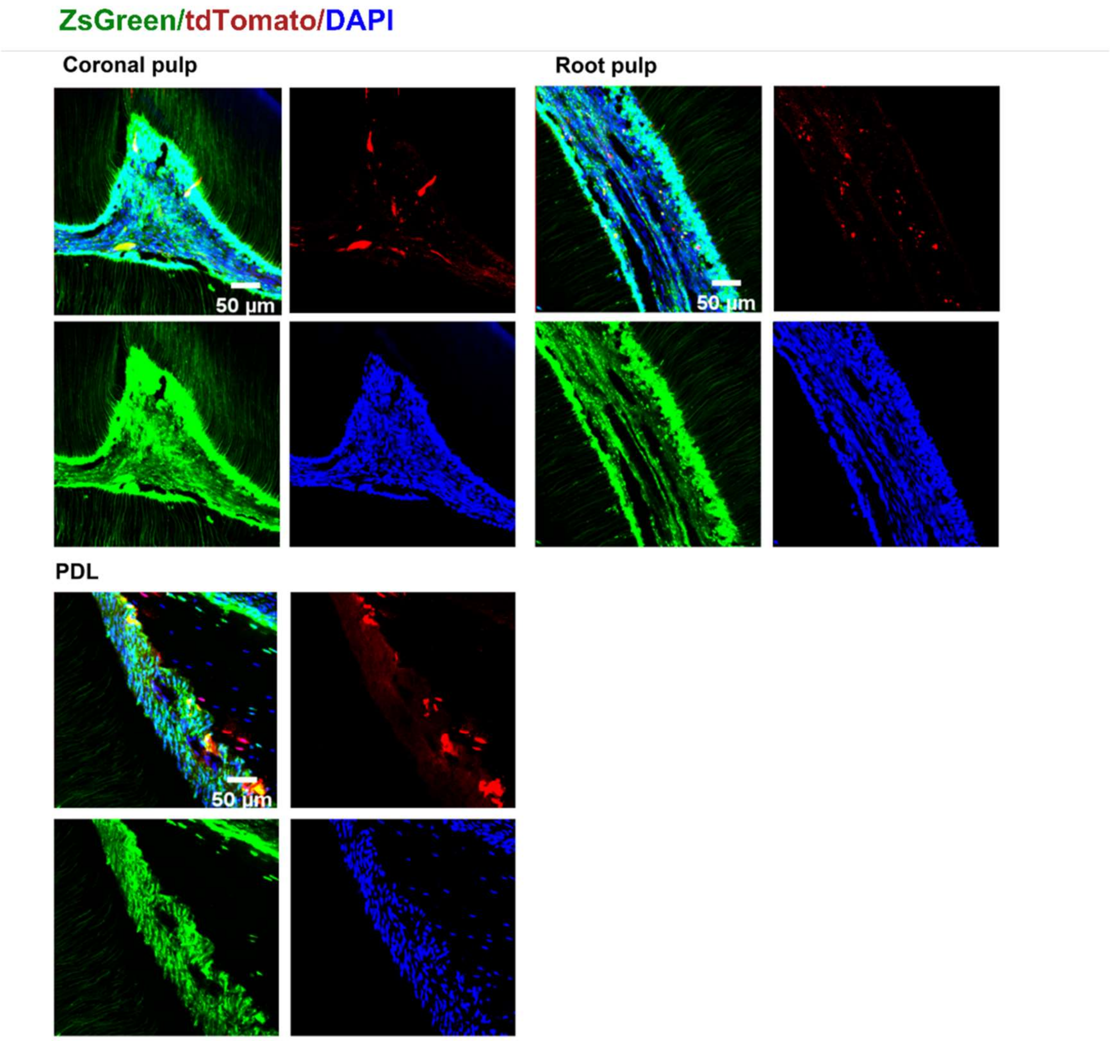


**Figure S11.** Representative images of coronal pulp, root pulp, and PDL of maxilla M1 of *Pdgfr-α*$^{CreER}$; *Nfatc1*$^{DreER}$; *LGRT* mice sample (tracing). The images were acquired by confocal microscope (scale bar = 50 µm).

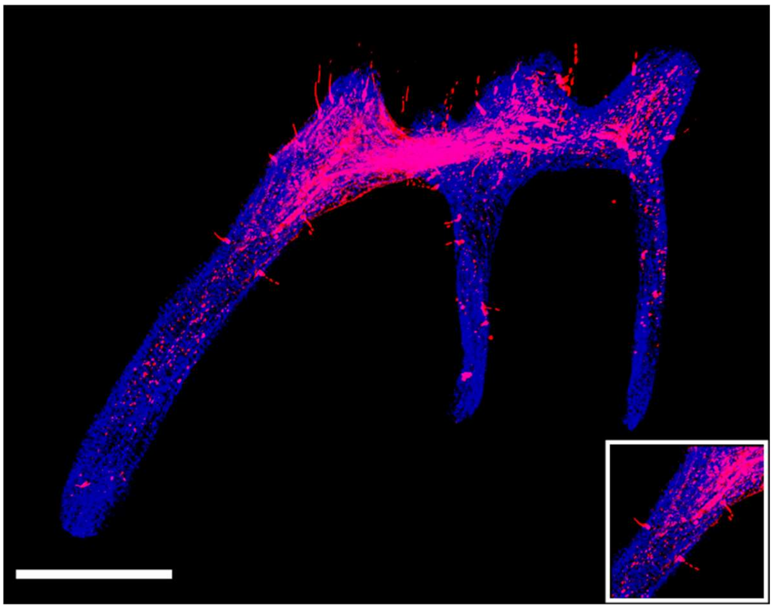

**Figure S12.** The tdTomato signal in pulp of maxilla M1 of *Pdgfr-α*$^{CreER}$*; Nfatc1*$^{DreER}$*; LGRT* mice sample (tracing) reconstructed by traditional serial section-based confocal imaging method (scale bar = 300 μm).

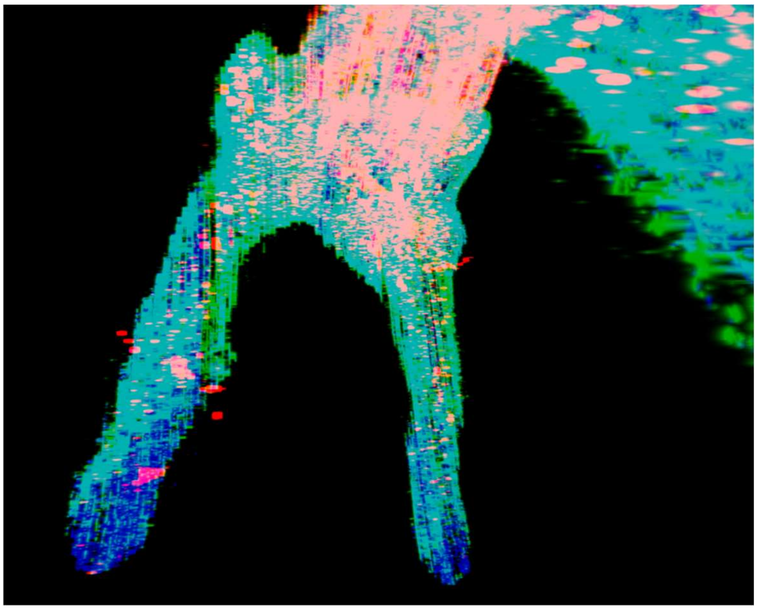

**Figure S13.** The discontinuities in the z-axis due to stratification of slices. The image was reconstructed by serial sections of maxilla M1 of *Pdgfr-α*$^{CreER}$*; Nfatc1*$^{DreER}$*; LGRT* mice sample (tracing) using Imaris. ZsGreen$^{+}$ cells in green, tdTomato$^{+}$ cells in red, DAPI in blue.

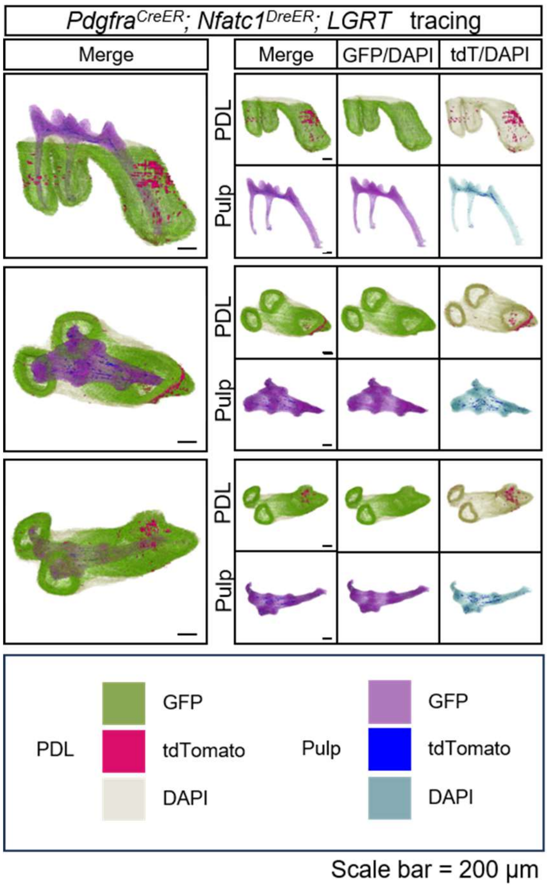

**Figure S14.** 3D reconstruction of maxilla M1 of *Pdgfr-α*$^{CreER}$; *Nfatc1*$^{DreER}$; *LGRT* mice (tracing) by DICOM-3D; PDL: ZsGreen$^{+}$ cells in green, tdTomato$^{+}$ cells in rose red; pulp: ZsGreen$^{+}$ cells in purple, tdTomato$^{+}$ cells in blue. The image stack was also displayed in buccal view, coronal view, and radicular view of pulp and PDL, respectively.

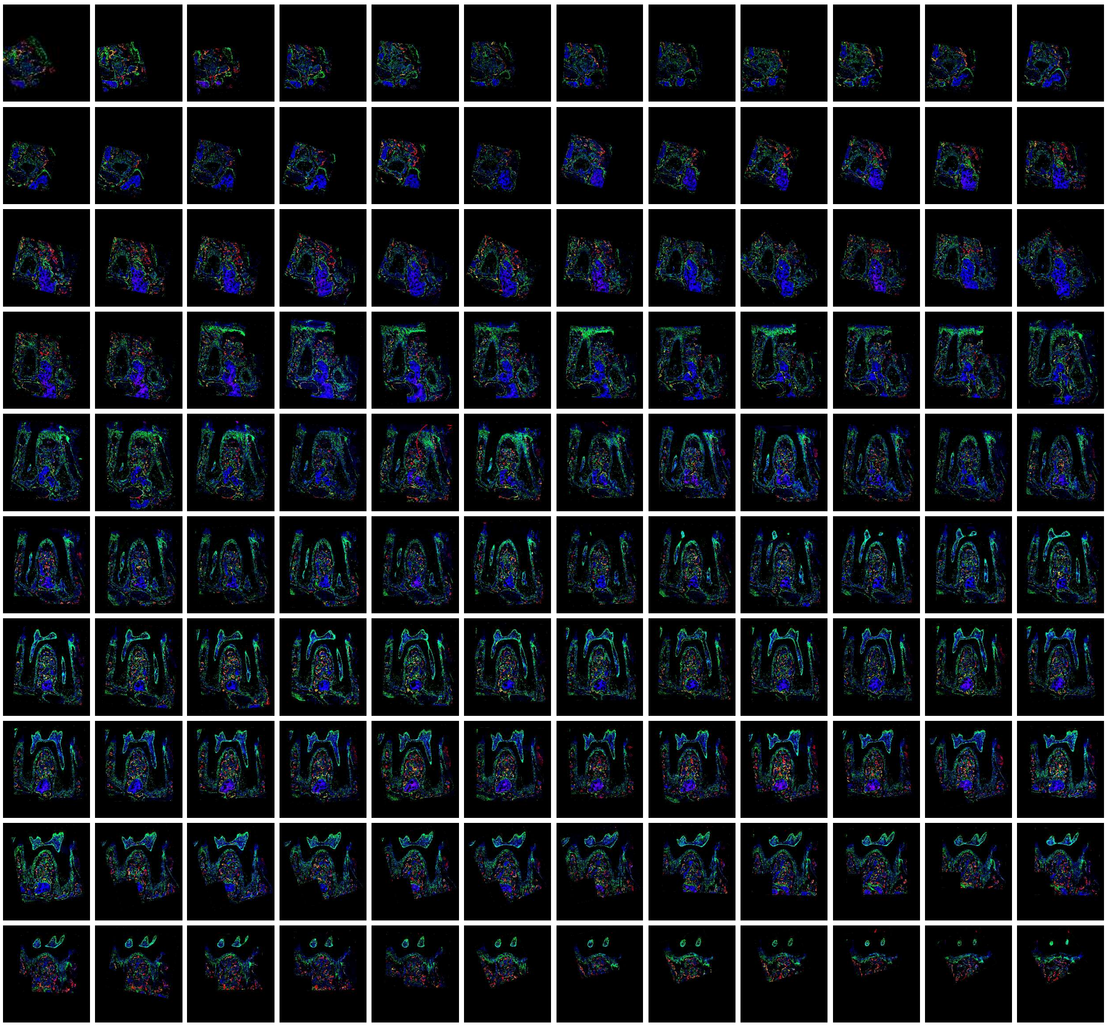

**Figure S15.** All consecutive slices (a total of 120 slices) for imaging of mandible M1 of *Pdgfr-α*$^{CreER}$; *Nfatc1*$^{DreER}$; *LGRT* mice (tracing 11 days). The images were acquired by confocal microscope, ZsGreen$^{+}$ cells in green, tdTomato$^{+}$ cells in red, DAPI in blue.

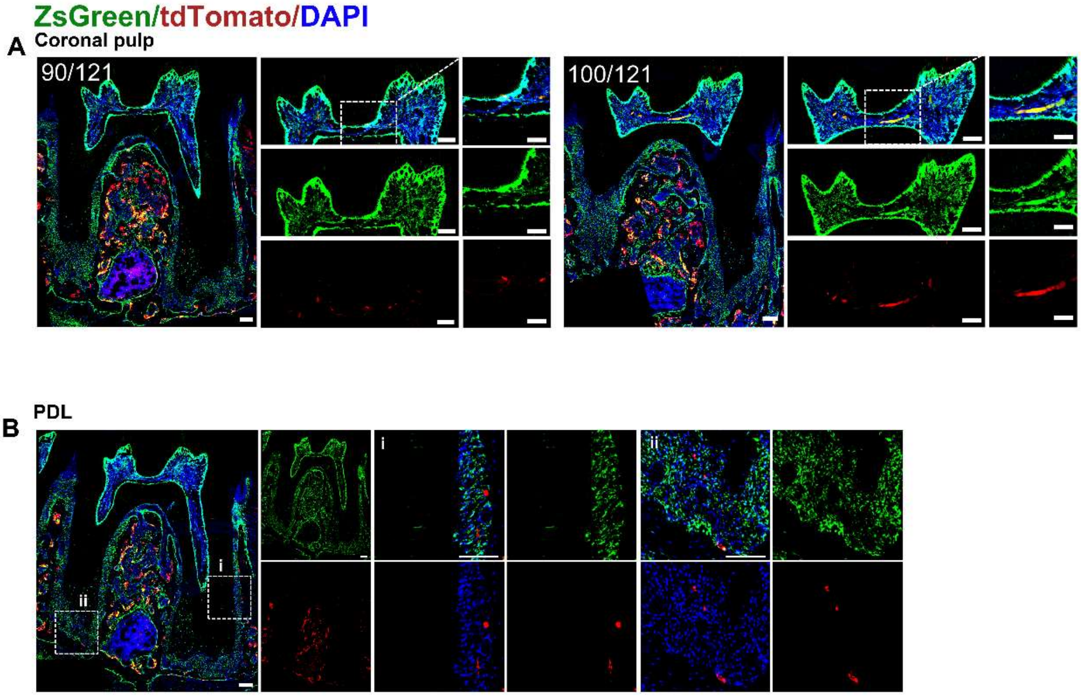


**Figure S16.** Representative images of coronal pulp (A) and PDL (B) acquired by confocal microscope of mandible M1 of *Pdgfr-α*$^{CreER}$*; Nfatc1*$^{DreER}$*; LGRT* mice sample (tracing). Scale bar = 100 μm.

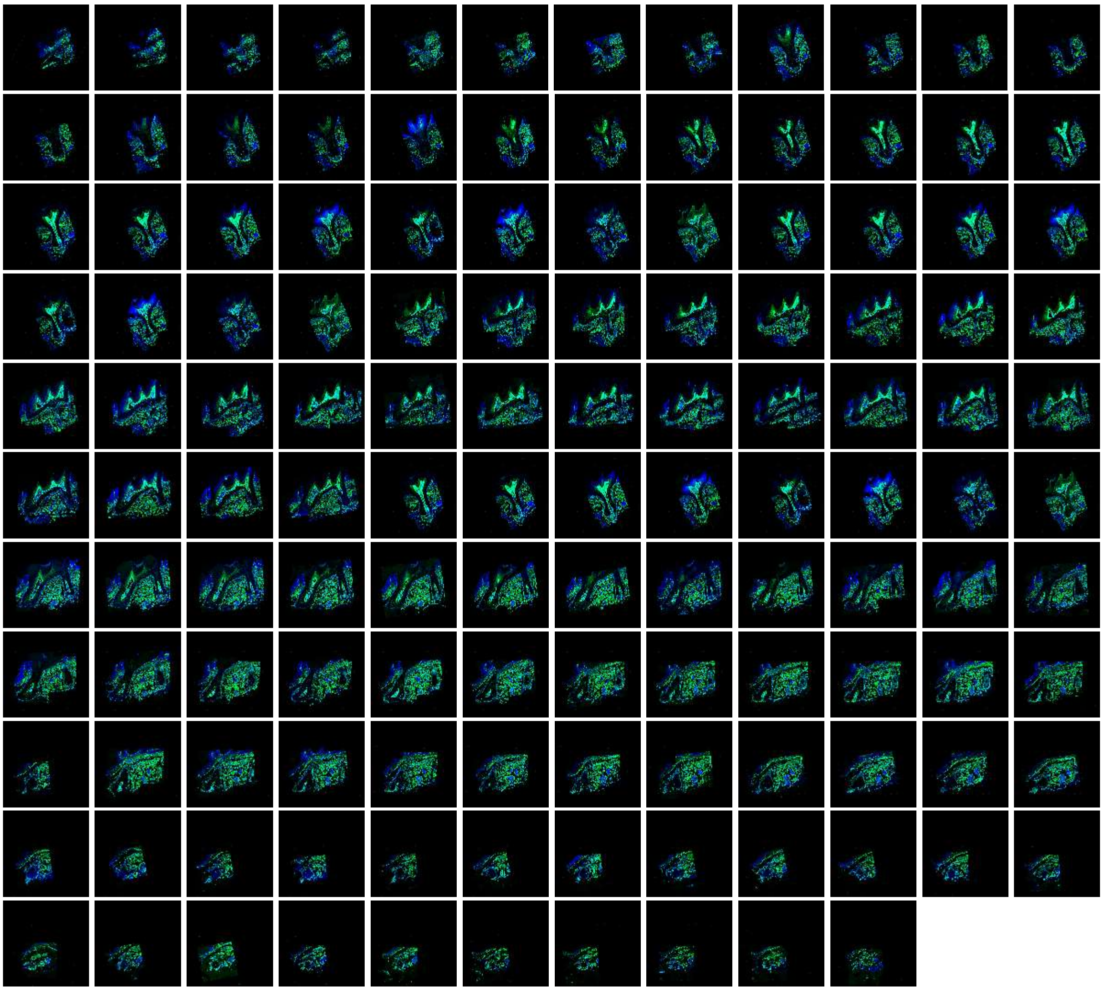

**Figure S17.** All consecutive slices (a total of 130 slices) for imaging of maxilla of *Pdgfr-a*$^{CreER}$*; IR1* mice. The images were acquired by confocal microscope, ZsGreen$^{+}$ cells in green and DAPI in blue.

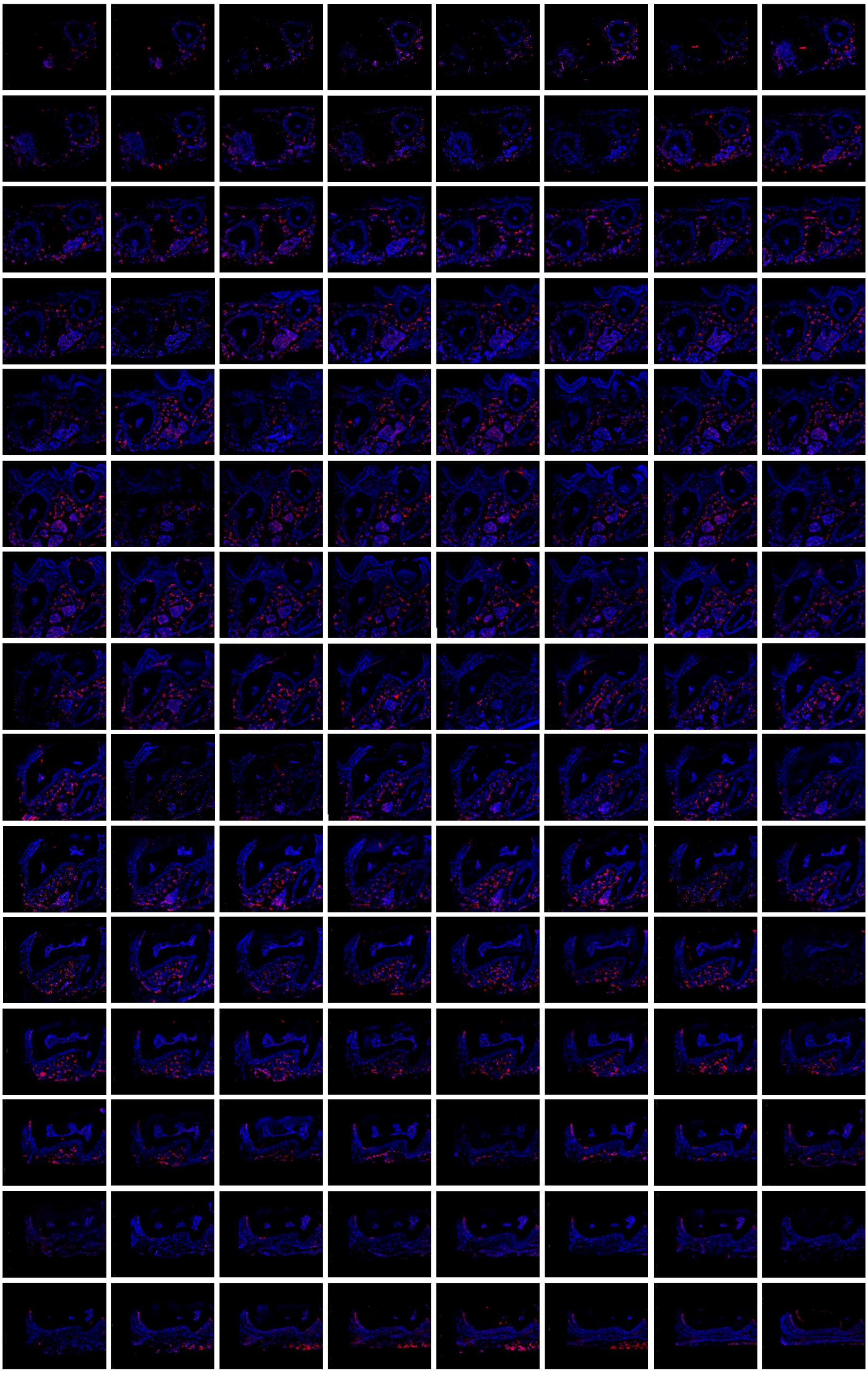

**Figure S18.** All consecutive slices (a total of 121 slices) for imaging of maxilla of *Nfatc1*$^{DreER}$*; IR1* mice. The images were acquired by confocal microscope, tdTomato$^{+}$ cells in red and DAPI in blue.

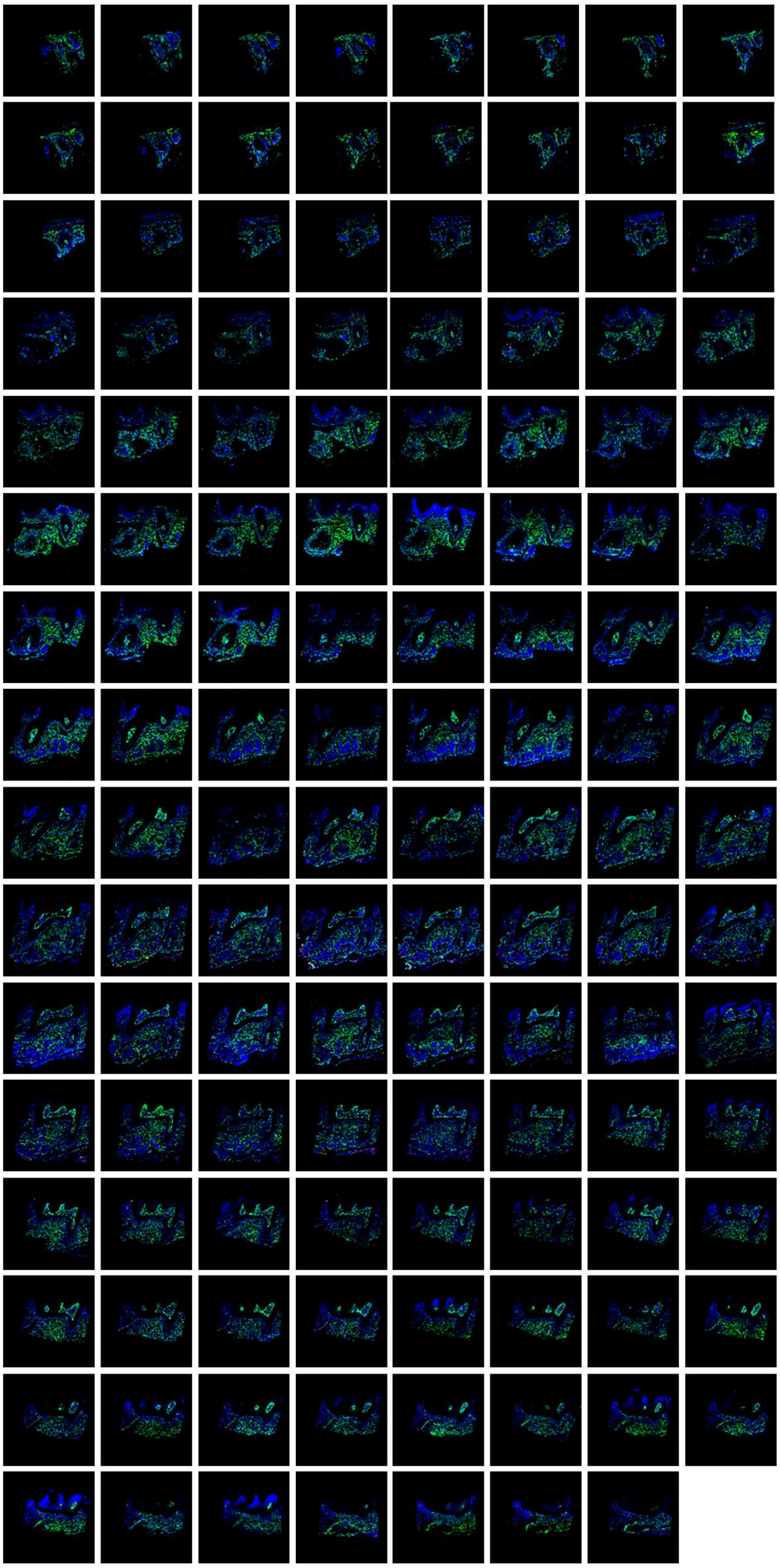

**Figure S19.** All consecutive slices (a total of 127 slices) for imaging of maxilla of *Pdgfr-a*$^{CreER}$; *Nfatc1*$^{DreER}$; *IR1* mice sample. The images were acquired by confocal microscope, ZsGreen$^{+}$ cells in green, tdTomato$^{+}$ cells in red, DAPI in blue.

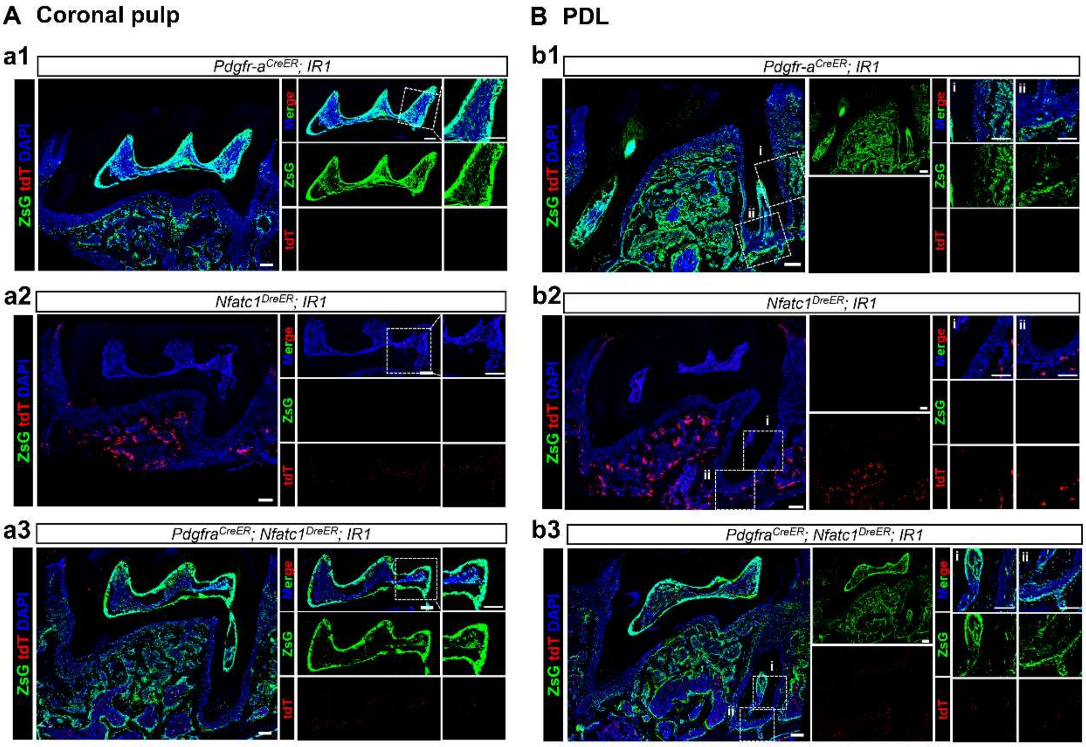


**Figure S20.** Representative images of coronal pulp (A) and PDL (B) acquired by confocal microscope of mandible M1 of *Pdgfr-a*$^{CreER}$*; IR1* (a1, b1), *Nfatc1*$^{DreER}$*; IR1* (a2, b2) and *Pdgfr-α*$^{CreER}$*; Nfatc1*$^{DreER}$*; IR1* (a3, b3) mice. Scale bar = 100 µm.

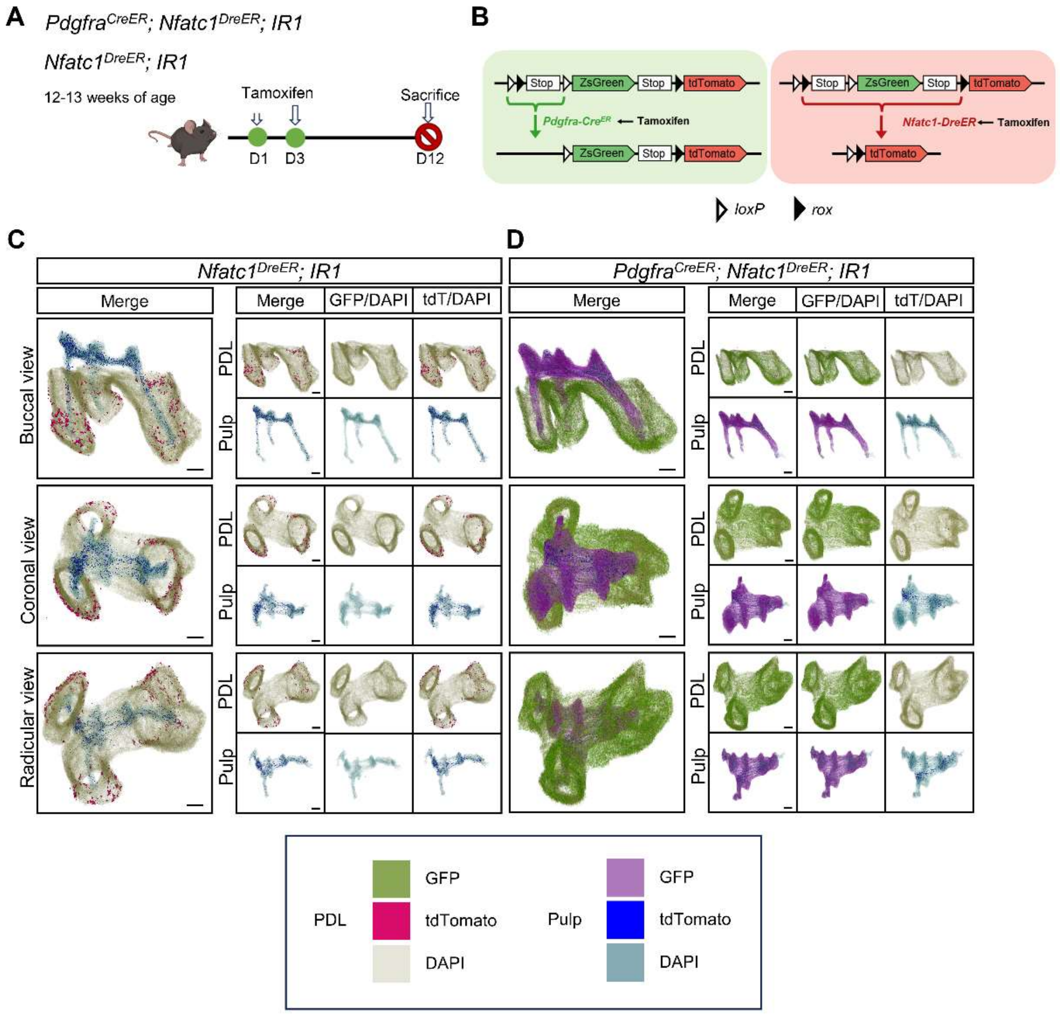


**Figure S21.** 3D reconstruction of maxilla M1 by DICOM-3D of maxilla M1 of *Nfatc1*$^{DreER}$*; IR1* and *Pdgfr-a*$^{CreER}$*; Nfatc1*$^{DreER}$*; IR1* mice; in PDL, ZsGreen$^{+}$ cells in green, tdTomato$^{+}$ cells in rose red; in pulp, ZsGreen$^{+}$ cells in purple, tdTomato$^{+}$ cells in blue. The image stack was displayed in buccal view, coronal view, and radicular view. Scale bar: 200 μm.

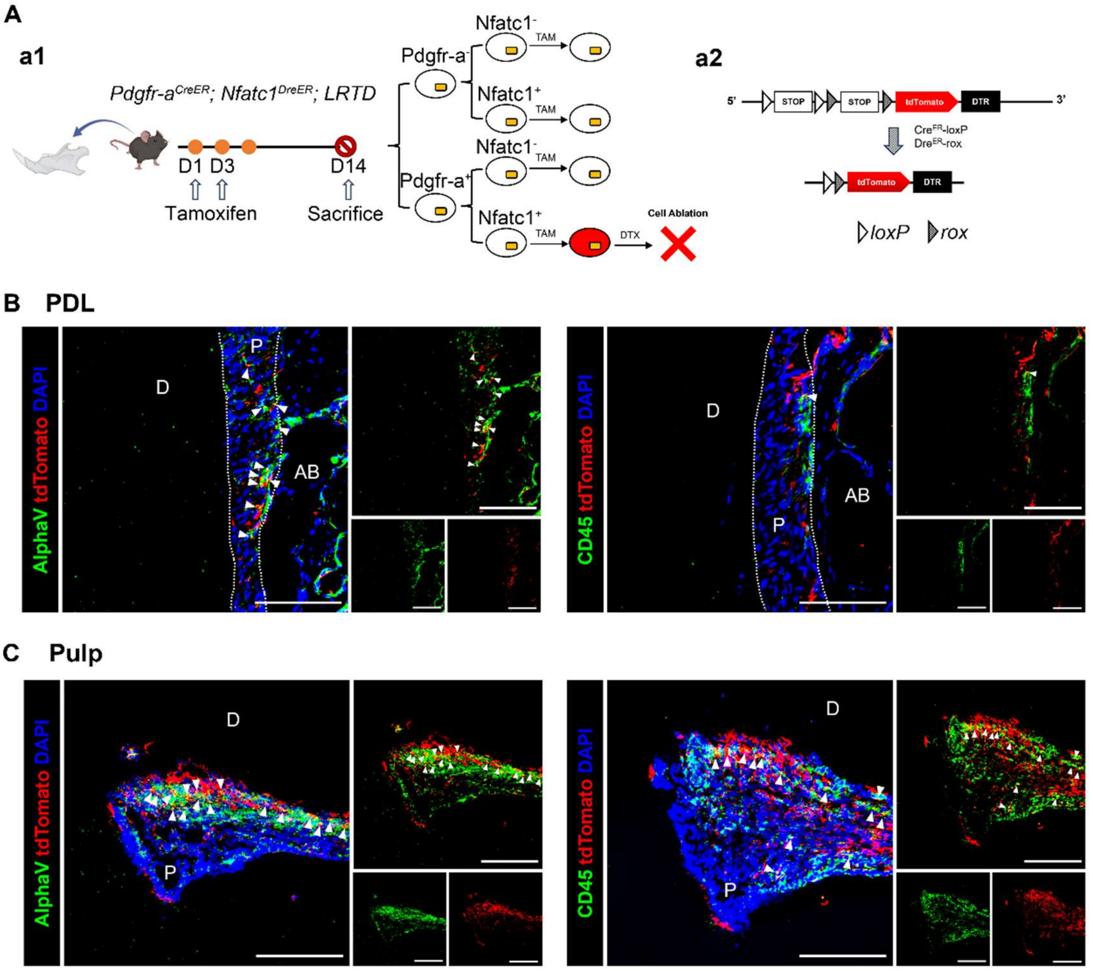


**Figure S22.** The identification of the types of PDGFR-$\alpha^+$ & NFATc1$^+$ cells (red) of dental pulp and periodontal ligament tissues. (A) Schematic illustration of lineaging tracing in *Pdgfr-α*$^{CreER}$*; Nfatc1*$^{DreER}$*; LRTD* mice. The mice were administrated with tamoxifen at D1 and D3, and sacrificed at D14. (B, C) Representative IF images of PDL B) and pulp C) of mandible M1 in P *Pdgfr-α*$^{CreER}$*; Nfatc1*$^{DreER}$*; LRTD* mice showing the MSCs marker AlphaV (left) and hematopoietic marker CD45 (right). Arrows B, C) indicate the co-localization of AlphaV/CD45 and tdTomato. D: dentin; AB: alveolar bone. P: PDL B), pulp C). Scale bar: 100 μm.

### Legends of supplemental videos

Video S1. Panoptic multicolor imaging of PDGFR-$\alpha^+$ cells(green) & NFATc1$^+$ cells(red) in the pulp and PDL area of maxilla M1 of *Pdgfr-α*$^{CreER}$*; Nfatc1*$^{DreER}$*; LGRT* mice (pulse), the whole-tissue imaging was achieved through TC procedure, related to Figure 2C.

Video S2. Panoptic multicolor imaging of PDGFR-$\alpha^+$ cells & NFATc1$^+$ cells in the pulp and PDL area of maxilla M1 of *Pdgfr-α*$^{CreER}$*; Nfatc1*$^{DreER}$*; LGRT* mice (pulse), the whole-tissue imaging was reconstructed from serial sections, related to Figure 3C.

PDGFR-α$^+$ cells in green, NFATc1$^+$ cells in red and DAPI in blue

Video S3. Panoptic multicolor imaging of PDGFR-α$^+$ cells & NFATc1$^+$ cells in the pulp and PDL area of mandible M1 of *Pdgfr-α$^{CreER}$; Nfatc1$^{DreER}$; LGRT* mice (pulse), the whole-tissue imaging was reconstructed from serial sections, related to Figure 4C. PDGFR-α$^+$ cells in green, NFATc1$^+$ cells in red and DAPI in blue

Video S4. Panoptic multicolor imaging of ZsGreen$^+$ cells (green) & tdTomato$^+$ cells (red) in the pulp and PDL area of maxilla M1 of *Pdgfr-α$^{CreER}$; Nfatc1$^{DreER}$; LGRT* mice (tracing 11 days), the whole-tissue imaging was achieved through TC procedure, related to Figure 5B.

Video S5. Panoptic multicolor imaging of ZsGreen$^+$ cells & tdTomato$^+$ cells in the pulp and PDL area of maxilla M1 of *Pdgfr-α$^{CreER}$; Nfatc1$^{DreER}$; LGRT* mice (tracing 11 days), the whole-tissue imaging was reconstructed from serial sections, related to Figure 6C. ZsGreen$^+$ cells in green, tdTomato$^+$ cells in red and DAPI in blue

Video S6. Panoptic multicolor imaging of ZsGreen$^+$ cells (green) & tdTomato$^+$ cells (red) in the pulp and PDL area mandible M1 of *Pdgfr-α$^{CreER}$; Nfatc1$^{DreER}$; LGRT* mice (tracing 11 days), the whole-tissue imaging was reconstructed from serial sections, related to Figure 7C. ZsGreen$^+$ cells in green, tdTomato$^+$ cells in red and DAPI in blue

Video S7. Panoptic multicolor imaging of PDGFR-α$^+$ cells (green) & NFATc1$^+$ cells (red) in cranium of *Pdgfr-α$^{CreER}$; Nfatc1$^{DreER}$; LGRT* mice (pulse), the whole-tissue imaging was achieved through TC procedure.

Video S8. Panoptic multicolor imaging of PDGFR-α$^+$ cells (green) & NFATc1$^+$ cells (red) in cranial sagittal suture of *Pdgfr-α$^{CreER}$; Nfatc1$^{DreER}$; LGRT* mice (pulse), the whole-tissue imaging was achieved through TC procedure.

Video S9. Panoptic multicolor imaging of PDGFR-α$^+$ cells (green) & NFATc1$^+$ cells (red) in cranial coronal suture of *Pdgfr-α$^{CreER}$; Nfatc1$^{DreER}$; LGRT* mice (pulse), the whole-tissue imaging was achieved through TC procedure.

Video S10. Panoptic multicolor imaging of ZsGreen$^+$ cells in the pulp and PDL area of maxilla M1 of *Pdgfr-α$^{CreER}$; IR1* mice, the whole-tissue imaging was reconstructed from serial sections, related to Figure 8. ZsGreen$^+$ cells in green and DAPI in blue.

Video S11. Panoptic multicolor imaging of tdTomato$^+$ cells in the pulp and PDL area of maxilla M1 of *NFATc1$^{DreER}$; IR1* mice, the whole-tissue imaging was reconstructed from serial sections, related to Figure 8. tdTomato$^+$ cells in red and DAPI in blue.

Video S12. Panoptic multicolor imaging of ZsGreen$^+$ cells & tdTomato$^+$ cells in the pulp and PDL area of maxilla M1 of *Pdgfr-α$^{CreER}$; NFATc1$^{DreER}$; IR1* mice, the whole-tissue imaging was reconstructed from serial sections, related to Figure 8. ZsGreen$^{+}$ cells in green, tdTomato$^{+}$ cells in red and DAPI in blue.
